# Refractoriness-Based Masking Yields Threshold-Specific Human Auditory Nerve Responses

**DOI:** 10.64898/2026.09.19.752910

**Authors:** Feifan Chen, Thomas J Stoll, Gerilyn R Jones, Ross K Maddox

## Abstract

Distinct auditory nerve (AN) fiber populations with varying thresholds and vulnerabilities to synaptopathy are observable in animal models, but standard non-invasive human measures lack the resolution to differentiate these groups of fibers. This fundamental limitation hinders the translation of findings in animals to human auditory pathologies. To overcome this, we developed a novel refractoriness-masking paradigm to non-invasively isolate threshold-specific AN fiber responses in humans. By presenting rapid click trains with cumulatively rising levels (45–105 dB peSPL) at a 250 μs inter-click interval, we sequentially masked lower-threshold fibers. Using a tympanic membrane electrode, we recorded compound action potentials (CAPs) in adults with normal hearing and isolated threshold-specific responses via waveform subtraction. Our results demonstrated that the isolated CAPs exhibited decreased latencies with increasing levels and a non-linear amplitude growth, plateauing at 90 dB peSPL before sharply increasing at 105 dB peSPL. This pattern reflects the sequential recruitment of AN fiber that were not effectively recruited by the preceding clicks. Validate conditions, in which the final two clicks had equal levels, produced substantially smaller CAPs, consistent with response suppression by refractoriness. Overall, this paradigm establishes a human measurement approach for characterizing level-threshold-dependent AN responses, providing a potential way to better investigate different AN pathologies, such as cochlear synaptopathy or auditory neuropathy.

## Introduction

The auditory nerve (AN) serves as the critical link between the cochlea and the central auditory system, transmitting the output of cochlear hair cells to the brainstem (Spoendlin & Schrott, 1989). Each type I AN fiber receives input from a single inner hair cell (Liberman, 1980) and is responsible for encoding various aspects of sounds, including intensity, frequency, and timing (Rutherford et al., 2021). While the response patterns of individual AN fibers can be directly recorded in animals through invasive recordings, electrophysiological measures available in human research, such as auditory brainstem responses (ABRs) or electrocochleography (ECochG), lack sufficient resolution to differentiate contributions from specific fiber populations. This limitation hinders further investigation of the role of specific groups of AN fibers in human sound coding or their pathology in neural degeneration. Here we sought to develop a novel paradigm based on neural refractoriness to better differentiate threshold-specific AN fibers and their electrophysiological responses in humans.

The most common categorization of AN fibers in mammals is high-, medium-, and low-spontaneous rate (SR) fibers (Liberman, 1978). These subtypes show distinct characteristics in their thresholds, dynamic ranges, temporal precision, and susceptibility to damage (Alamri & Jennings, 2023; Verhulst et al., 2018). Invasive animal recordings have shown that high-SR fibers dominate the population (over 50%) and are highly sensitive to low-level sounds, whereas low-SR fibers are less sensitive but contribute to encoding suprathreshold stimuli and have been hypothesized to be critical for sound perception in noise (Kujawa & Liberman, 2015). Recent animal studies have shown that low-SR fibers are more vulnerable to degeneration than other fibers due to aging or noise exposure (Fernandez et al., 2020; Kujawa & Liberman, 2009, 2015; Sergeyenko et al., 2013; Valero et al., 2017), leading to loss of synapses known as cochlear synaptopathy. However, results of neurophysiological (e.g., wave I of the ABR) and/or behavioral measures (e.g., speech perception in noise) in human studies have not found a reliable metric for cochlear synaptopathy or correlation with speech in noise performance (N. Bramhall et al., 2019; N. F. Bramhall et al., 2017; Grose et al., 2019; Guest et al., 2018, 2019; Mehraei et al., 2016; Sergeyenko et al., 2013; Suresh & Krishnan, 2020). Apart from the heterogeneity of study designs, species differences in response properties of AN fibers populations may explain the inconclusive results from human data. The correlation between response thresholds of AN fibers and their SR has been observed in non-primate animal models (e.g., cats, gerbils, guinea pigs), but this relationship did not hold in the few studies of non-human primate models that examined this correlation (Joris et al., 2011; Nomoto et al., 1964). Due to this species difference, we focus on threshold-specific responses in this study, rather than attempting to distinguish responses by spontaneous rate.

While direct recordings from individual AN fibers are not feasible in humans, the summed response of AN activity can be measured through ECochG and ABR. ECochG records electrical potentials generated from the cochlea and AN fibers (Ruth et al., 1988), whereas ABR captures the evoked electrical activity generated of the auditory brainstem and midbrain (Eggermont, 2019). The activity of AN fibers can be reflected by the compound action potential (CAP) in ECochG recordings or ABR wave I (Jewett & Williston, 1971; Pienkowski et al., 2018). Notably, both techniques capture the activity from a collection of AN fibers, which synchronize to stimuli but vary in location, spontaneous rate, and susceptibility to adaptation (Heil & Peterson, 2015). Therefore, it is not possible to differentiate the contribution of different AN fiber types to the summed potential with current methods.

To overcome this limitation, we developed a paradigm to sequentially mask out the response from AN fibers with lower response thresholds. The paradigm relies on the refractoriness of AN fibers (Miller et al., 2001), which is an instant decrease in discharge probability (i.e., refractory period) following a spike (Brown et al., 1996; Hodgkin & Huxley, 1952). As a universal property of spiking neurons, fibers that respond to the preceding sounds will be refractory at a short inter-stimulus interval (ISI) and less likely to be activated by the following stimuli. Consequently, the refractoriness of AN fibers could serve as a potential tool to separate fiber responses by manipulating the level of stimuli and the interval between them.

Here, we used brief click trains with rising levels and short ISIs to isolate AN populations by threshold in humans, measured via ABR wave I and the CAP. We hypothesized that 1) preceding lower-level clicks in the train will selectively suppress lower-threshold AN fibers such that they cannot respond to subsequent clicks due to refractoriness, 2) the neural response to the final, highest-level click in the train will be dominated by the high-threshold AN fibers that are responsive but not yet saturated, and 3) the characteristics of neural responses (e.g., ABR wave I or CAP) will thus reflect sequential recruitment of AN populations with different response thresholds. This method to distinguish response patterns across different AN fibers will not only help to bridge the gap between human and animal research in auditory nerve functions but may also have potential as a more sensitive proxy for neural degeneration in the auditory periphery, such as auditory neuropathy or cochlear synaptopathy.

## Methods

### Participants

Twenty participants (age = 28.7±6.2, 10 males, 10 females) were recruited from the community. Exclusion criteria included any known history of sensorineural or conductive hearing loss, hearing disorder, acoustic neuroma, neuropathy, neurological disorders or claustrophobia. All participants underwent pure tone audiometry (AC40, Interacoustics) where hearing thresholds in each ear at 250, 500, 1000, 2000, 4000, and 8000 Hz were obtained. ER-3 insert earphones were used for testing. All participants were screened for hearing thresholds ≤ 20 dB HL, except for 2 participants who showed mild hearing loss at 8 kHz (<35 dB HL). All participants gave their written, informed consent in compliance with the University of Michigan Medical School Institutional Review Board (IRBMED HUM00268049). Participants were compensated for their time.

### Electrophysiology Recording

#### Stimuli and Experiment Design

We designed a refractoriness-masking paradigm to isolate fiber threshold-specific responses in humans. The paradigm utilizes rapid click trains with cumulatively rising sound levels. By setting an inter-click interval (ICI) within the absolute refractory period of the auditory nerve, the preceding, lower-level clicks activate lower-threshold AN fibers and keep them in a refractory state. Consequently, the neural response to the final, highest-level click in a train should be dominated by the higher-threshold fibers that have not yet been activated. We then isolate the final-click responses by subtracting the response to a shorter click train (containing only the preceding masking clicks) from the response to the target click train.

#### Experimental session

The clicks in each train were 100 μs pulses. The ICI in each train was first set as 1 ms but substantial neural activity from lower-threshold fibers was observed in subsequent click responses, indicating incomplete masking (see Results for more details). Therefore, we shortened the ICI to 250 μs, as the absolute refractory period of human AN fibers ranges from 200 μs to 600 μs (Morsnowski et al., 2006; Skidmore et al., 2022). These stimuli consisted of five rapid click trains with one, two, three, four, or five clicks and an absent stimulus condition. Each click was 15 dB higher in level than the one that preceded it, such that the clicks ranged from 45 to 105 dB peSPL in the train with five clicks (**Fig. 1A i-iii**). The preceding lower-level clicks within a train were presumed to serve as the maskers of the response to the next click, but only for fibers that were responsive to that lower level. Null stimulus trials (∅) where no sounds were presented were included to measure and allow us to remove the activity from the previous set of clicks.

**Fig. 1.**
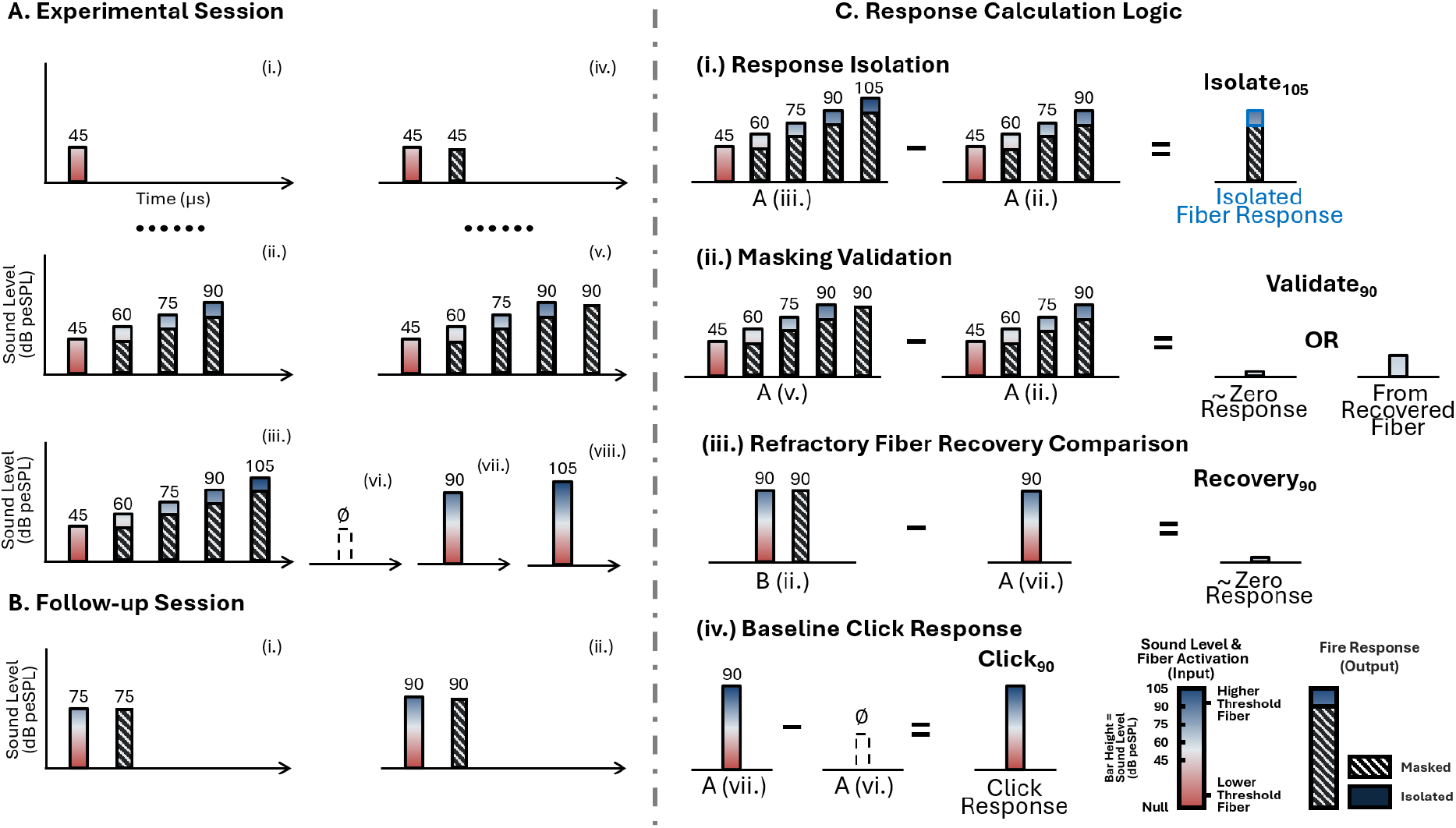
Refractoriness masking paradigm. (A) Experimental Session has 5 click trains with cumulatively rising levels (e.g., i-iii), 4 click trains where the last two clicks have identical sound levels (e.g., iv, v), 2 single-click references (vii, viii), and a null stimulus (vi). (B) Follow-up Control Session includes 2 trains with cumulatively rising levels, 2 trains with the last two clicks identical, 2 single clicks and 2 paired-clicks (i-ii) to evaluate the potential fiber recovery from neural refractoriness. (C) Response condition calculation logic demonstrates how to compute *Isolate*, *Validate*, *Recovery* and *Click* responses mathematically through waveform subtraction (e.g., subtracting a shorter preceding train from the target train to isolate the final click).

In order to verify the effectiveness of refractoriness masking, we included four additional click trains in which the level of each click in each train was 15 dB higher than the preceding one, except for the last two clicks that were identical in sound levels (**Fig. 1A iv, v**). The levels were from 45 dB to 90 dB peSPL. We reasoned that if refractory masking were effective, there would be no response to the final click.

Finally, two single-click stimuli where the sound level was 90 or 105 dB peSPL (**Fig. 1A vii, viii**) were included as a reference for the response morphology and magnitude for each participant.

In total, the experimental session included two sub-sessions with an ICI of either 1 ms or 250 μs. All twenty participants finished 1 ms sub-session first and we invited them back for the 250 μs sub-session. Among them, thirteen participants underwent the second sub-session. For each sub-session, twelve stimuli were included: 5 trains for isolating responses, 4 trains for verifying effectiveness, 2 single clicks, and 1 null stimulus. Each stimulus condition had 15,000 trials with 40 ms between trial starts. Stimuli were delivered in alternating polarity. All conditions were mixed and pseudo-randomized by creating balanced randomization of transitions between different conditions where the probabilities of each transition are identical. Stimuli were presented diotically via ER-3 earphones at 48,000 Hz using custom Python scripts and open-source library expyfun (Larson et al., 2014).

#### Follow-up control session

In the 250 μs sub-session, while there were limited CAP waves in the *Validate* responses across all levels, we observed a peak component at 75 and 90 dB peSPL that resembled a typical component following the CAP (see **Fig. 3A**). This lack of complete masking could have been from previously active neurons coming out of their absolute refractory period and firing again, or a result of some neurons that could have spiked in response to the previous click not doing so (as firing is probabilistic, especially near threshold) and then responding to the next one. We designed a follow-up control experiment to distinguish which mechanism was at play. Four out of thirteen participants were invited back for the session. During the session, there were two click trains for isolating fiber responses (up to 70 or 90 dB peSPL), two trains for verifying effectiveness (up to 70 or 90 dB peSPL), and two single-click stimuli at 70 and 90 dB peSPL. Notably, we also added two paired-click stimuli, in which the click trains only included two clicks with identical levels (either 70 or 90 dB peSPL) (**Fig. 1B i, ii**). This resulted in a total of 8 stimulus conditions. Each condition had 22,500 trials with a duration of 40 ms. All clicks were presented with an ICI of 250 μs and pseudo-randomized as in the experimental session.

#### Auditory Brainstem Response (ABR)

Participants were seated in reclining chairs in a dark sound-treated and electrically-shielded booth (IAC, North Aurora, IL, USA). ABRs were recorded using a BrainVision ActiCHamp Plus EEG system and two EP-Preamp ABR preamplifiers (BrainVision, LLC, Greenboro, SC, USA). Two bipolar channels were used to record the ABR, with the noninverting electrode placed at FCz and the inverting channels placed on the left and right earlobes. The ground electrode was placed on the high forehead (Fpz). EEG was recorded with a sampling frequency of 10,000 Hz. To minimize impedances between the electrodes and scalp and improve signal quality, dead skin and oils were removed by scrubbing the earlobes and forehead with NuPrep and an alcohol pad, with a target impedance of ≤5 kΩ.

#### Electrocochleogram (ECochG)

ECochG was recorded by a tympanic membrane (TM) electrode developed by Simpson et al. (2020). Additional modifications in the process for making TM electrodes were described in a previous study (Stoll et al., 2025). The tip of the TM electrode was coated with Signa electrode gel (Parker Laboratories, Fairfield, NJ). Then it was set on the central location of the right eardrum by an experienced clinician using Bebird ear tweezers with a live camera (Heifeng Zhizao, Shenzhen, China) and fixed in place by a standard foam insert. The tweezers were modified by putting thin heat shrink on the jaws, increasing grip and making it easier to manipulate the fine wire of the electrodes. As with the ABR electrodes, the TM electrode was plugged into an EP-preamp to be recorded in a bipolar montage, with the TM electrode used as the non-inverting electrode and the ipsilateral earlobe as the inverting electrode. In addition to visual inspection and participant feedback to confirm that the electrode was on the eardrum, responses to a brief (three minute) recording of randomly-timed (Polonenko & Maddox, 2019) 90 dB peSPL clicks (mean rate: 20 stim/s) were examined. Impedances for these TM electrodes are typically quite high, so correct electrode placement and recording fidelity was verified by the presence of a clear CAP prior to starting the experiment. Our results verified that the TM electrode is a reliable measure of AN physiology and is more sensitive relative to ABR wave I (**Fig. 2**), which is consistent with previous studies that utilized the TM electrode (Goodman et al., 2023; Simpson et al., 2020).

**Fig. 2.**
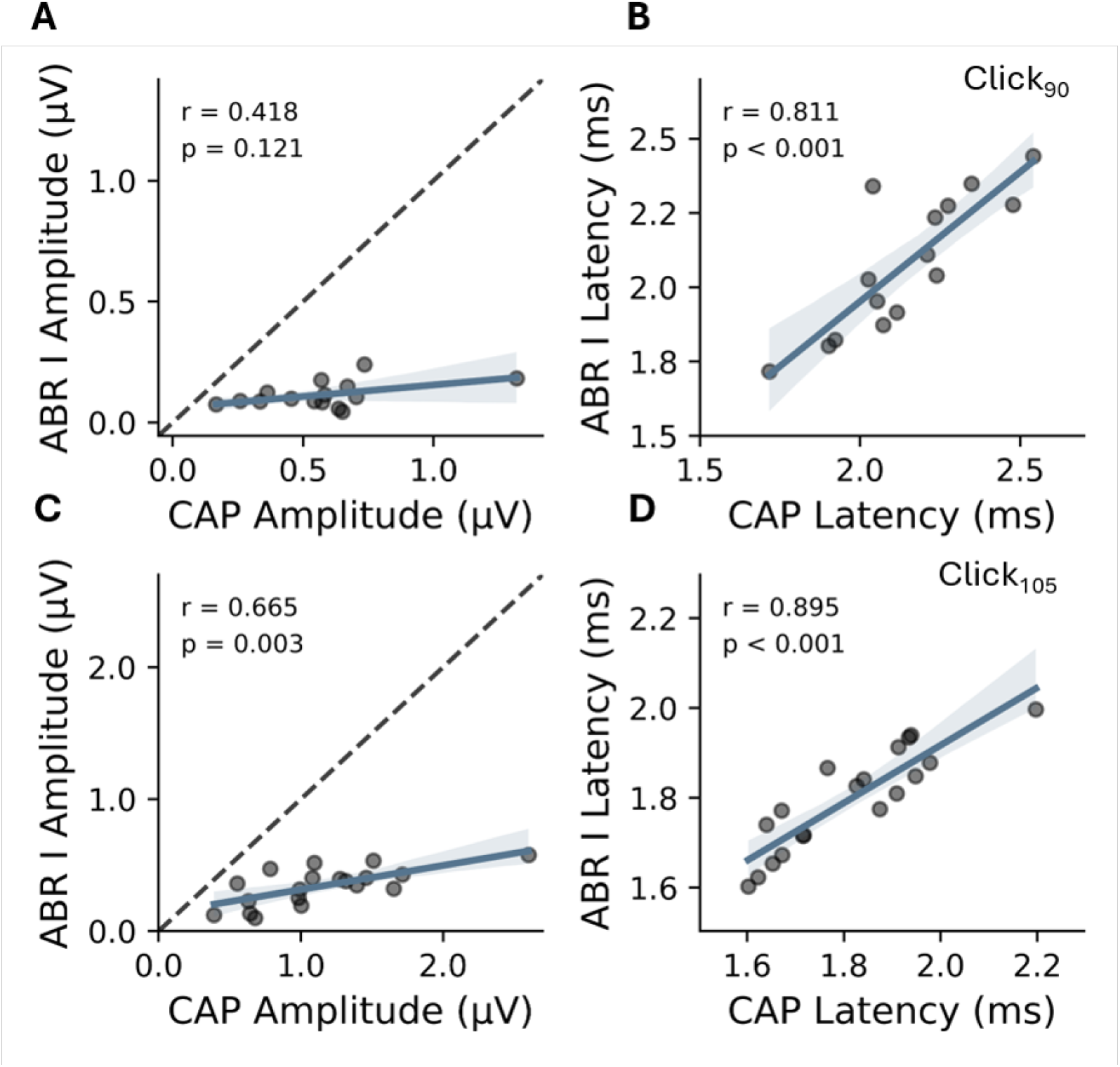
TM electrode as a reliable measure of AN physiology. Correlation analyses between CAP and ABR wave I responses to single-click stimuli at 90 (A & B) and 105 dB peSPL (C & D). Two neural responses were recorded simultaneously. Significant correlations were observed for response latencies at both sound levels and response amplitude at 105 dB peSPL. Compared to ABR wave I amplitude, CAP amplitudes were larger for both sound levels, supporting the enhanced sensitivity and reliability of the TM electrode over traditional scalp electrodes.

### Electrophysiology Analysis

#### Preprocessing

Data were preprocessed in Python using the MNE package (Gramfort et al., 2014). First, the raw data were filtered by first-order causal infinite impulse response (IIR) bandpass filters, which were 30–2000 Hz for ABR and 150–2500 Hz for ECochG. A causal IIR notch filter at odd multiples of 60 Hz up to 540 Hz were applied to remove power line noise. Data were then epoched from −20 ms to 40 ms relative to the onset of the trial after accounting for the ∼1 ms delay introduced by the acoustic tubing.

#### Bayesian Weighting

After preprocessing, a Bayesian weighting method was used to improve SNR when calculating the average response. Specifically, the variance of the noise was estimated for each trial from the pre-stimulus baseline window (−20 to −2 ms relative to onset) and the trial weight was set as the inverse of this variance. To further mitigate the influence of transient artifacts, an outlier rejection step was applied: trials with weights falling outside 1.5 times the interquartile range of the weight distribution were excluded. The weights were then normalized such that they summed to one across trials for each electrode and each condition. Finally, the Bayesian-weighted average responses were determined through summing weighted responses.

#### Waveform Subtraction

Given the short ICI, the response to the final click in each train inevitably overlapped with the residual responses to the preceding clicks. To isolate the target response to the final click, we computed *Isolate* waveforms by subtracting the response to the click train that was shorter by one click (i.e., containing only the preceding clicks) from that of the target click train (**Fig. 1Ci**). To validate the effectiveness of neural refractoriness, *Validate* waveforms were calculated by the response difference between the trains with the identical last two clicks and the trains containing only the preceding clicks (**Fig. 1C ii**). To test the fiber recovery from refractoriness in the follow-up control session, *Recovery* waveforms were calculated by subtracting the corresponding single-click from the paired-click train (**Fig. 1C iii**). For the *Click* waveforms, the baseline activity to the null stimulus (i.e., ∅) was subtracted from click-evoked responses to get rid of the average response from the prior trial.

To account for differences in onset timing of the final click across conditions, response waveforms were temporally realigned to enable consistent comparison of the stimulus-evoked responses across conditions. Specifically, the difference waveforms were shifted such that the zero point on the time axis corresponded to the onset of the final click for all conditions. The trailing portion of the signal was zero-padded to preserve the original epoch length, although this region was not included in any analyses.

#### ABR Analysis

ABR wave V for each response was picked automatically by taking the maximum and minimum points of the waveform between 6 and 13 ms. Accuracy of the automatic peak picking was confirmed through visual inspection, with manual adjustments when necessary. Wave V amplitudes were defined by the difference between the peak and following trough voltages, whereas wave V latencies were defined as the time of the selected peak voltage.

#### CAP Analysis

The compound action potential (CAP) was similarly picked automatically by taking the minimum points waveform between 1 and 5 ms. CAP amplitudes were defined as the magnitude of the trough voltage and CAP latencies were defined as the time of the selected trough voltage. The ratios of CAP amplitude between two consecutive target response conditions were calculated to minimize the variability due to non-sensory factors.

#### RMS Analysis

Root mean square (RMS) values for CAP responses were calculated as an index of overall neural recruitment, which reflects neural activity throughout the entire analysis window, regardless of their precise temporal alignment. The time range for deriving RMS was 0.5–7.5 ms after the onset of final click in each train, which enabled the inclusion of the full CAP waveform at all levels. Assuming that the noise component and the evoked neural signals are statistically independent, response magnitudes were quantified using a noise-corrected RMS algorithm. To account for the contribution of background noise to the recorded signal, a signal-cancellation technique was applied using the same Bayesian weighting mentioned above. Trials were first ranked in ascending order of their weights. The polarity of trial data was then inverted for every second trial in the ranked sequence (i.e., alternating polarity based on weight rank). A weighted average of these polarity-inverted trials was computed to cancel out the deterministic response, effectively isolating the residual background noise. The noise-adjusted RMS amplitude was subsequently calculated by subtracting the mean square of this noise estimate from the mean square of the response (equation 1) and taking the square root while preserving the sign (equation 2),

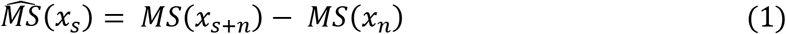

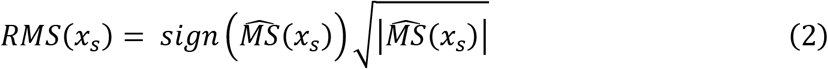

where MS indicates the mean-squared value, RMS the root-mean-squared value, *x_s_*_+*n*_ the average response (which contains signal and noise), and *x_n_* the noise estimate. Thus, a negative sign indicates larger noise than signal, or vice versa.

### Behavioral Measures

We included three behavioral tasks as indices of AN fiber function: an extended high frequency (EHF) test (Rieke et al., 2017), the Coordinate Response Measure (CRM) test (Gallun et al., 2013), and the amplitude modulation (AM) detection test (Whiteford et al., 2017). All tasks were implemented using the *expyfun* Python library (Larson et al., 2014). Stimuli were presented at a sampling rate of 48,000 Hz via an RME Babyface Pro sound card and delivered through Sennheiser HD 650 Headphones. The details of each behavioral task and results are included in the supplementary information.

### Statistical Analysis

Statistical analysis was performed in Python using the Pingouin package (Vallat, 2018). For both the 1 ms and 250 μs sub-sessions, one-way repeated-measure ANOVAs were conducted for the amplitudes and latencies of the CAP and ABR wave V to examine the effect of masking on distinguishing the fiber-specific response pattern. The within-subject factor was sound level. The assumption of sphericity was assessed using Mauchly’s test. When the assumption of sphericity was violated, degrees of freedom were adjusted using the Greenhouse-Geisser correction. Significant main effects or interactions were followed up with pairwise comparisons. The Pearson correlation coefficient (*r*) was applied separately to investigate the correlation between different neural responses, and between the CAP ratio and behavioral measures, including the EHF, CRM and AM detection tests. The significance level was set at alpha = 0.05. The Benjamini-Hochberg procedure was applied for all tests to correct for multiple comparisons with a false discovery rate (FDR) of 5%.

To test the hypothesis that the morphology reconstructed from isolated-fiber responses was similar to corresponding single-click responses despite amplitude modulation, we conducted a waveform similarity analysis. For each participant, the vector of voltage values was extracted for both waveforms in specific time window: 0–3.1 ms for the 90 dB condition; 0–2.2 ms for 105 dB condition. We calculated the Pearson correlation coefficient between these vectors to quantify morphological adherence. Rather than relying on a parametric statistical test, we employed a bootstrapping procedure (10,000 resamples with replacement) on the *z*-transformed values to estimate the non-parametric 95% confidence interval (CI) of the group mean. The z-score CI was subsequently transformed back to Pearson’s *r* CI.

## Results

### Distinguishable fiber-specific responses at the inter-click interval of 250 μs

We first tested if preceding lower-level clicks in a click train could selectively suppress lower-threshold AN fibers from responding to subsequent clicks due to the refractoriness, which serves as the foundation of the paradigm. More specifically, the CAP to the *Validate* conditions was computed and compared with that of the *Isolate* conditions. If lower-threshold fibers were excited and remained in the refractory period when the following click was presented, *Validate* responses should be very small across all sound levels. This would confirm successful masking of the lower-threshold fibers by the preceding clicks and indicate the *Isolate* responses are dominated by higher-threshold AN fibers that did not respond to the lower-level click stimuli.

**Fig. 3.**
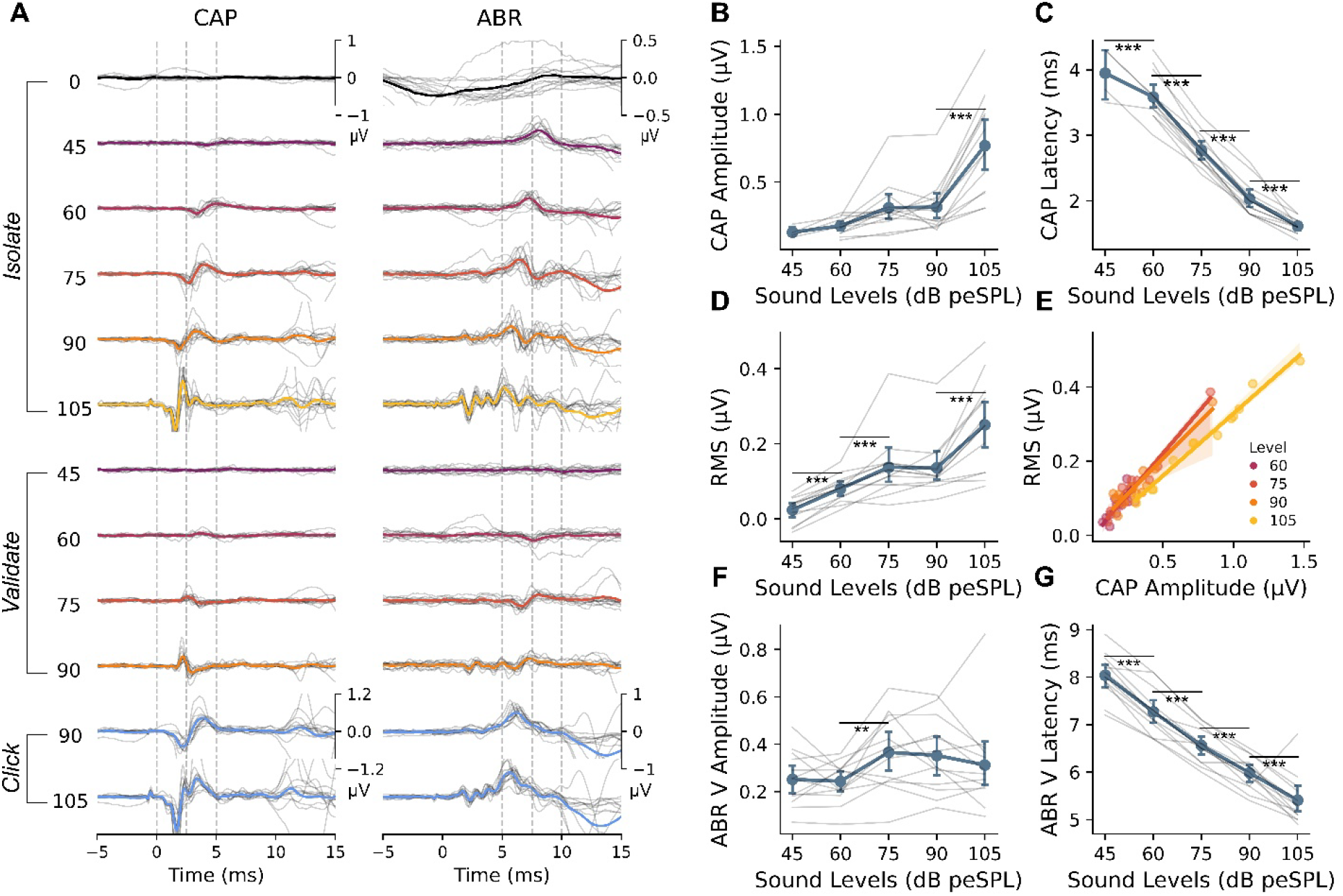
Grand average CAP and ABR waveforms in the 250 μs experimental sub-session. (A) Grand average waveforms across 3 conditions (*Isolate*, *Validate*, *Click*), showing robust *Isolate* CAPs starting from 60 dB peSPL and predominantly inhibited AN responses in the *Validate* conditions. (B-C) *Isolate* CAP amplitudes and latencies demonstrated significant level effects, characterized by decreasing latencies and a sharp amplitude increase at 105 dB peSPL. (D-E) Noise-adjusted RMS amplitudes of *Isolate* responses and their strong, significant correlation with CAP peak amplitudes across all measured sound levels. (F-G) The change of ABR wave V amplitudes and latencies over the sound level.

The grand average CAP and ABR responses are plotted in **Fig. 3A**. As expected, the CAP to each *Validate* condition was largely inhibited regardless of stimulus level. By contrast, we observed robust *Isolate* responses starting from 60 dB peSPL. There was a main level effect on both CAP amplitude (*F (4,12) = 33.616, p < .001, η²_p_ = .794*) (**Fig. 3B**) and latency *F (4,12) = 119.485, p < .001, η²_p_ = .943*) (**Fig. 3C**) across individual sound levels. We observed significantly shorter latencies as sound level increased (*p < .05* for all pairwise comparisons), whereas the increase in mean amplitude was not significant between 45 and 90 dB peSPL, followed by a significant and sharp increase at 105 dB peSPL (*p < .05*). For both *Isolate* and *Validate* responses at 45 and 60 dB peSPL, a CAP was sometimes not observed, presumably due to the lower sound levels.

Since previous studies proposed RMS as a potential biomarker for AN fiber physiology (Fujihira et al., 2024, 2026), we also measured the RMS of the *Isolate* response with the window ranging from 0.5 to 7.5 ms after the onset of the target click (**Fig. 3D**). The RMS showed a similar level effect as the CAP amplitude (main level effect: *F (4,48) = 39.011, p < .001, η²p = .507*), which was also supported by the strong correlation between CAP amplitude and RMS at all levels (60dB: *r = .841, p < .001*; 75dB: *r = .965, p < .001*; 90 dB: *r = .946, p < .001*; 105 dB*: r = .991, p < .001*) (**Fig. 3E**). The pairwise comparisons showed an increase of RMS as level went up, except from 75 dB to 90 dB peSPL. The RMS quantifies the total neural output and overall fiber recruitment across the analysis window, whereas peak amplitude is highly driven by neural synchrony. Therefore, the significant correlation suggested sufficient neural synchrony of recruited fibers at each sound level, reflecting the intact sound encoding of AN fibers in young adults with normal hearing. Comparisons of ABR parameters responses across levels are shown in **Fig. 3F&G** and reported in **Table 1**.

**Table 1.** ANOVA of ABR parameters at the inter-click interval of 250 μs.

| Parameters | Source | df | F | p-value | $\eta^2_p$ |
| --- | --- | --- | --- | --- | --- |
| Amplitude | Level | 4 | 3.571 | <.05* | .116 |
|  | Residual | 48 |  |  |  |
| Latency | Level | 4 | 188.4 | <.001*** | .977 |
|  | Residual | 48 |  |  |  |

Note: \*: $p < .05$ , \*\*\*: $p < .001$

To further verify the *Isolate* responses represent distinct neural populations, we compared the *Click* responses with the summed *Isolate* responses up to the corresponding level (**Fig. 4A&B**). We hypothesized that the waveform morphology of the two responses should be preserved if the *Isolate* responses represent threshold-specific fiber responses. For both the 90 dB and 105 dB conditions, overall similar waveforms and strong correlations were observed between the summed *Isolate* and *Click* responses (90 dB: mean *r* = .91, 95% CI [.87, .95]; 105 dB: mean *r* = .89, 95% CI [.87, .91]).

**Fig. 4.**
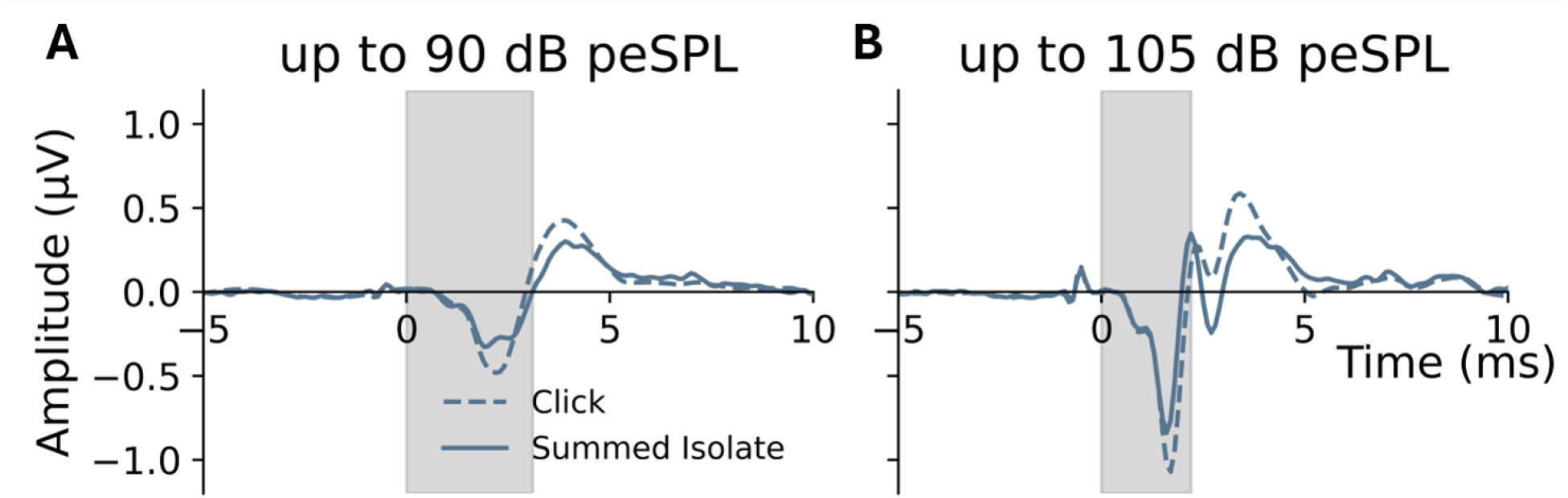
Isolated threshold-specific CAP responses could reconstruct single-click evoked responses. Morphological comparisons between single-click responses and the summed *Isolate* responses up to 90 dB peSPL (A) and 105 dB peSPL (B). The significant waveform correlation marked in grey (mean r = .91 for 90 dB; mean r = .89 for 105 dB) validated that the temporal morphology of the neural representation is conserved independent of amplitude, confirming the validity of the response isolation.

An exploratory analysis was conducted to investigate the relationship between threshold-specific fiber responses and behavioral measures. There was no significant correlation between any neural measures and any behavioral measures in participants with normal hearing (**Supplementary Fig. 2&3**). We note, however, that the participants were younger adults with normal hearing thresholds and no noise exposure history was collected. Thus, we do not have sufficient data to explore the parameter space, and no strong conclusions can be drawn from the absence of significant correlations.

### Recovery from refractoriness does not account for peak components in *Validate* **responses**

Although the CAP was greatly diminished in the *Validate* condition, a response component was present in the 75 and 90 dB peSPL conditions, and the amplitude of this component increased with sound level. One potential explanation of peak components in the *Validate* responses could be attributed to the activity of lower-threshold fibers that were excited by early low-level clicks but recovered from refractoriness during the interval between the second-to-last and last clicks. Thus, we invited 4 participants back for a control follow-up session. In this session, the ICI was again 250 μs. We introduced two new conditions with click trains that only included two clicks with identical levels (**Fig. 1B i,ii**). Through this paradigm, we aimed to maximally excite the AN fibers that are responsive up to these sound levels with the first click, greatly reducing the likelihood of fibers responding to the second click. We computed the *Recovery* waveforms by subtracting single-click responses from paired-click responses at the same level to isolate the response to the second of the paired clicks. We hypothesized that if the peak component of *Validate* responses at high levels is driven by a group of AN fibers that were excited by the preceding lower-level clicks that had recovered from refractoriness, we should not observe the same response component in the *Recovery* responses at those levels.

As shown in **Fig. 5A&B**, at both 75 and 90 dB peSPL levels, *Validate* and *Recovery* conditions showed similar grand-average CAP morphology, suggesting that the response component observed in the *Validate* condition was not driven by fibers that had recovered from refractoriness. Instead, it likely stems from the fact that neural responses are probabilistic and some with thresholds below the click level (and especially below but near) did not respond to the first click but did to the second.

**Fig. 5.**
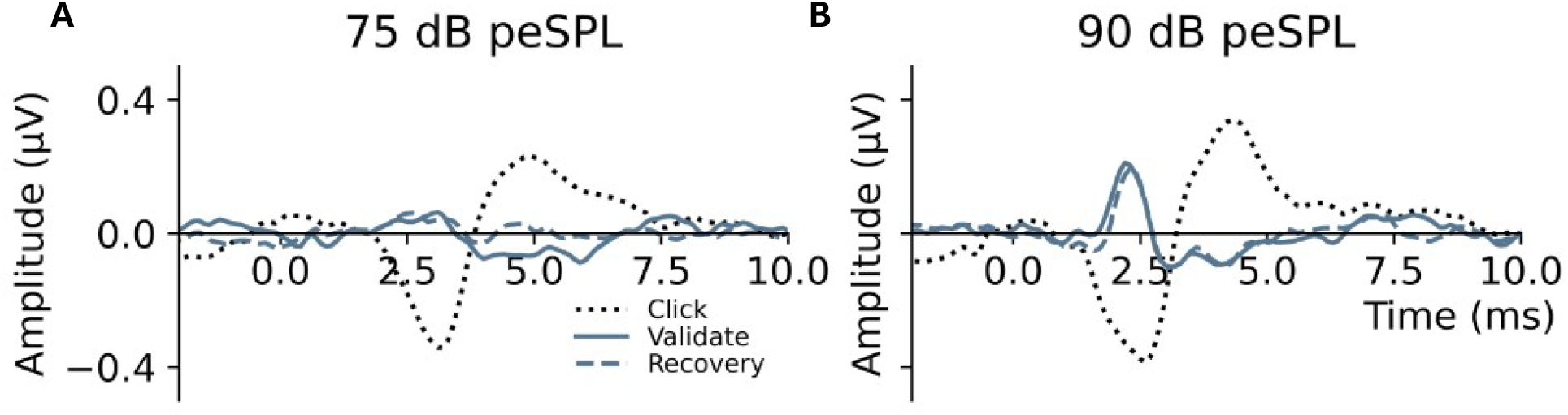
Identical CAP morphology between the *Validate* and *Recovery* conditions. Grand average waveforms from the follow-up control session comparing single-click, *Validate*, *Recovery* conditions at 75 dB peSPL (A) and 90 dB peSPL (B). The identical CAP morphology observed between the *Validate* and *Recovery* conditions suggested that the peak components seen in high-level *Validate* responses are not driven by the activity of lower-threshold fibers recovering from refractoriness.

## Discussion

This study aimed to develop and validate a novel refractoriness-masking paradigm to disentangle the threshold-specific contributions of human AN fiber populations. We utilized rapid click trains with cumulatively increasing intensities (from 45 to 105 dB peSPL) and inter-click interval of 250 μs to sequentially suppress activity elicited by lower-level clicks. We could then isolate the activity from AN fibers recruited only at higher sound levels through systematic waveform subtraction. By employing a high-fidelity electrode directly on the tympanic membrane, we demonstrated that this approach can effectively capture nuanced neural responses more reliably than is possible through traditional ABR measures (i.e., wave I).

Critically, the observed amplitude increase and latency decrease of *Isolate* responses support successful isolation of fibers due to sequential fiber recruitment from lower-threshold to higher-threshold populations with increasing sound levels (**Fig. 3B&C**). Further, the strongly reduced *Validate* responses support effective suppression of responses to a repeated same-level click at 250 μs, validating the effectiveness of this masking paradigm (**Fig. 3A**). Together, these threshold-specific electrophysiological patterns hold potential to serve as sensitive, non-invasive proxies for assessing human cochlear neural degeneration of different fiber populations, such as cochlear synaptopathy.

### *Isolate* Response: threshold-specific fiber recruitment

The main finding of our paradigm was the ability to isolate threshold-specific responses. One pattern we observed was the non-linear amplitude change as a function of level: it exhibited an ascending slope below 75 dB peSPL but reached a plateau at 90 dB peSPL and followed by an increase at 105 dB peSPL. This aligned with the observed input-output curve of the single-click CAP response in previous human studies (Alamri & Jennings, 2023; Ohashi et al., 2005; Salomon & Elberling, 1971; Yoshie, 1968; Zöllner et al., 1976) that showed the response size grew slowly at low intensity and then rapidly at high intensity, with the inflection point around 50 dB SL (≈ 85 dB peSPL). These findings are consistent with the statement that there are two populations of sensory units that dominate the amplitude growth (Yoshie, 1968). The nonlinear amplitude growth may reflect recruitment of fibers with higher response thresholds. Notably, high-level clicks may also cause the spread of excitation in cochlear regions, which could jointly contribute to the marked amplitude increase at 105 dB peSPL. Increasing click level from 75 to 95 dB peSPL shifts the predominant cochlear contribution to the CAP from more apical regions (approximately 2–4 kHz) toward more basal regions (approximately 8–20 kHz) (Alamri & Jennings, 2023; Elberling, 1974). Thus, the pronounced amplitude increase in the *Isolate_105_* condition may reflect both recruitment of higher-threshold fibers and recruitment of fibers from additional cochlear regions that were weakly or not activated by the preceding 90-dB click and therefore remained available to respond.

The reconstruction of *Click* CAP morphology by summing *Isolate* responses further supported that the derived components capture threshold-specific contributions to the click-evoked CAP. Using neural adaptation to separate AN-fiber responses has been investigated in numerous human studies by using the paired-click paradigm (i.e., a two-click train with short ICI) (Bidelman & Syed Khaja, 2014; Fujihira et al., 2024, 2026; Lee et al., 2021; Murnane et al., 1998; Ohashi et al., 2005). Our paradigm showed several advantages: all previous studies applied relatively long ICI, with the shortest ICI being 0.7 ms (Bidelman & Syed Khaja, 2014). Given that the absolute refractory period of human AN fibers is between 200 μs and 600 μs (Morsnowski et al., 2006; Skidmore et al., 2022), an ICI longer than 0.5 ms might not be sufficient to keep responding fibers in the refractory period, which was consistent with the incomplete masking seen in our 1 ms ICI experiment (see **Supplementary Fig. 1**). Therefore, our paradigm exhibited lager and more effective masking effect on fibers that have been recruited. Furthermore, in our paradigm, we applied multiple clicks with cumulative sound levels in a train, which kept the evoked responses temporally aligned with the onset of each click. Compared to the paired-click paradigm, it substantially reduced the overlapping contribution of preceding clicks, yielding a difference waveform dominated by activity associated with the final click. Also, the isolated responses evoked by a wide range of sound levels provide more information of the AN electrophysiology than the paired-click paradigm that only employed a single level. Given diverse physiological functions of different fiber populations, this might serve as a potential tool to investigate their roles in different temporal and spatial encodings.

It should be noted that we attribute the source of the isolated CAP responses to “threshold-specific” rather than “SR-specific” AN fibers. At relatively short ICIs, the decreased responses to the last click were hypothesized to be dominated by lower-SR fibers in numerous human studies (Fujihira et al., 2024, 2026; Murnane et al., 1998; Ohashi et al., 2005). Although various non-primate animal models have demonstrated that low-SR fibers exhibit higher response thresholds compared to their high-SR counterparts (Relkin et al., 1995), primates appear to possess distinct fiber characteristics. Specifically, two previous studies have shown no correlation between AN response thresholds and SR in primate models (Joris et al., 2011; Nomoto et al., 1964). Consequently, the direct translation of non-primate SR-threshold relationships to humans should be approached with caution. Thus, when responses are isolated based on neural response threshold, as in the present study, a relationship with SR should not be assumed.

### Amplitude and RMS: a more comprehensive measure of AN physiology

Peak amplitude measures depend upon neural synchrony – large populations of neurons firing at the same time. Due to this sensitivity to synchronous firing, amplitude alone might not be a sufficient or fully reliable metric, because it is vulnerable to non-pathological factors such as anatomical differences and response spread/smearing induced by temporal dispersion. By contrast, RMS could serve as a complementary metric for measuring AN physiology, since RMS reflects the overall neural recruitment and the cumulative volume of neural discharge within a specified analysis time window. Compared to peak amplitudes, RMS is inherently less sensitive to small jitters in neural firing patterns. This can be supported by our results that while RMS and amplitude showed similar trends as a function of level, the increase from 45 to 60 dB and from 60 to 75 dB was only significant for RMS values (**Fig. 3D**).

The advantage of including two complementary measures has been validated by recent evidence that demonstrated that RMS can capture degraded neural activity related to aging whereas peak amplitude measurements cannot (Fujihira et al., 2024). Further, a follow-up study that applied the paired-click paradigm also demonstrated that, compared to a control group, the noise-exposed group had significantly smaller RMS values of the second-click response but no significant group difference in the amplitude (Fujihira et al., 2026). Therefore, combining the response characteristics from these two parameters provides a clearer picture of the neural physiology of threshold-specific fibers. Further study is necessary to characterize the response pattern of threshold-specific fiber responses among people with higher risk of cochlear neural degenerations, such as age related neural degeneration (Dias et al., 2024; Fabrizio-Stover et al., 2026), which may result in degraded neural synchrony of AN fiber responses. While reduced AN response amplitudes have been reported in older participants (Fabrizio-Stover et al., 2026; Märcher-Rørsted et al., 2026), the effect of aging on amplitude and RMS values of threshold-specific responses has not been studied.

Although we did not observe any significant correlation between neural and behavioral measures among young adults, further investigation on clinical groups or those with higher risk of cochlear neural degeneration is needed to explore the sensitivity of neural measures for predicting temporal encoding and speech perception in noise. Specifically, as we isolated threshold-specific fiber activity, the amplitude and RMS slope or threshold-specific/single-click amplitude and RMS ratio could reduce the variance due to inter-subject variability and non-sensory factors (e.g., channel placement).

### Suppressed *Validate* Response: the effect of neural refractoriness

In AN fibers, refractoriness is subdivided into two sequential phases: the absolute and the relative refractory periods. The absolute refractory period refers to an interval where fibers are entirely incapable of generating a subsequent action potential, regardless of the magnitude or intensity of the next stimulus. By contrast, during the relative refractory periods, fibers can be elicited, but the threshold for activation is significantly elevated. The estimation for the absolute and relative refractory period remain consistent across studies, typically ranging from 200–600 μs ms and 0.6–1.4 ms (He et al., 2018; Morsnowski et al., 2006; Skidmore et al., 2022; Wiemes et al., 2016). Our results are also consistent with this estimate, as the *Validate* responses were mostly suppressed with the ICI of 250 μs but showed detectable CAP in the 1 ms sub-session (**Supplementary Fig. 1**), which lies in the relative refractory period. Similar electrophysiological results were observed in the paired-click paradigm where the neural responses were substantially, but not fully, suppressed with an ICI of 700 μs (Bidelman & Syed Khaja, 2014).

To date, the neural mechanism underlying the paired-click paradigm in most studies has been adaptation (Ohashi et al., 2005; Parham et al., 1996; Relkin & Doucet, 1991), which is a decrease in response sensitivity due to lessened neurotransmitter release at the synapse between the hair cell and AN fibers (Harris & Dallos, 1979; Smith, 1977). In contrast, the current paradigm leveraged the refractoriness of AN fibers that inhibits the discharge probability after a spike (Miller et al., 2001). The time scales of these two recovery processes (i.e., refractoriness and adaptation) vary significantly. For instance, human CAP and ABR showed a partial amplitude recovery from 1 ms, yet full recovery to an unadapted state required much longer intervals that are larger than 100 ms (Bidelman & Syed Khaja, 2014; Fujihira et al., 2024; Lasky, 1997; Ohashi et al., 2005). While direct comparisons between those studies and our study are not straightforward, due to the essential difference in neural mechanisms leveraged to mask fibers, the isolated CAP responses in the current study are also influenced by neural adaptation, but it is likely not the major factor. Ohashi et al. (2005) observed that the recovery of second-click CAP with the 50 ms ICI and 40 dB nHL (≈ 75 dB peSPL) was around 90%. Although the recovery of response decreased as the level increased, the amplitude of second-click response was still more than 75% of the first click at 60 dB nHL (≈ 95 dB peSPL). This was in contrast with the limited *Validate* responses in the current study at 90 dB peSPL. Taken together, given the experiment is 2-hour long with the inter-trial interval of 40 ms, the effect of neural adaptation is inevitable as previous studies suggested (Ohashi et al., 2005; Parham et al., 1996). However, within a click train refractoriness should impose greater influence on suppressing lower-level fibers and isolating threshold-specific responses.

Previous studies also took advantage of differences in recovery times of different human AN fiber populations to isolate fiber-specific responses (McClaskey et al., 2022; Murnane et al., 1998; Zeng et al., 1991). For instance, it has been proposed that when using a paired-click paradigm, long-term human CAP recovery reflects the contribution of low-SR (i.e., high-threshold) AN fibers (Murnane et al., 1998), which is consistent with the animal evidence (Relkin et al., 1995; Relkin & Doucet, 1991). Given the hypothesis that low-SR fibers are important for suprathreshold hearing and more vulnerable to neural degeneration, older adults might experience a disproportionate loss or deficit of low-SR fibers. This was supported by the finding that the older adults required less time for AN responses to recover from prior stimulation compared to young adults (McClaskey et al., 2022). Taken together, we suspect that older adults with normal hearing should have smaller Target CAP responses at higher levels but relatively intact responses at lower levels under this refractory masking paradigm. Future work will be needed to test this hypothesis.

Another observation in our data was a “peak” component in the *Validate* responses at 75 and 90 dB peSPL. The same peak component was observed in the *Recovery* responses (**Fig. 5**), suggesting they are not generated by the fibers that were excited by preceding lower-level clicks and recovered from refractoriness before the final click hit. While additional experiments would be needed to determine the source of this component, we suspect the stochastic nature of neuron activation might be the driving factor. Should the second-to-last click have a high probability to excite a given population of neurons, many of them will fire and enter the refractory period, but not all. The remaining neurons from this population would then fire at the time of the final click, resulting in the unexpected responses in the *Validate* condition. Alternatively, the component may reflect a nonlinear cochlear interaction between the final two equal-level clicks. At short ICIs, the basilar-membrane response to the second-to-last click may still be ongoing when the final click reaches the same cochlear region. The two mechanical responses can therefore overlap and interact (Charaziak et al., 2020). Specifically, the preceding response can alter the response evoked by the final click, and the final click can also alter the portion of the preceding response that continues after its arrival. This latter interaction is particularly relevant to our subtraction method, which assumes that the preceding-click response is identical in trains with and without the final click. If the final click suppresses part of the ongoing preceding response, that component will not be cancelled during subtraction. Instead, it will remain as a negative residual and may appear as a late, positive component. Nevertheless, this interpretation remains tentative because Charaziak et al. (2020) measured basilar-membrane motion rather than neural responses. It remains unclear whether this mechanical interaction is transmitted through inner-hair-cell transduction, synaptic release, and auditory-nerve spike generation to produce the component observed here.

### Limitations

There are several limitations that should be noted. First, the present paradigm uses broadband click stimuli that do not resolve the cochlear place from which threshold-specific neural responses originate. Particularly, as sound level increases responses receive contributions from broader and more basal cochlear regions (Alamri & Jennings, 2023; Elberling, 1974). Thus, the sharp increase in the *Isolate*_105_ condition may be attributed to both higher-threshold fibers and fibers from broader cochlear regions that were less effectively activated by the preceding clicks. The concurrent increase in RMS is consistent with greater overall neural recruitment but also cannot distinguish these threshold- and place-dependent contributions. Future studies combining refractoriness masking with frequency-specific masking, or computational modeling of cochlear excitation across characteristic frequencies, will provide greater insight into the place of origin of these high-level responses. Second, 250 μs was chosen to fall within the range of absolute refractory periods reported for electrically stimulated human AN fibers (Morsnowski et al., 2006; Skidmore et al., 2022). Here, we applied acoustic stimulation in which refractoriness cannot be completely separated from adaptation. Finally, the participants recruited were mostly young adults with normal hearing thresholds. While the exploratory analysis did not reveal the significant relationship between neural and behavioral responses, future studies that recruit participants with a wider age range will better sample the feature space and may shed light on the effect of aging on threshold-specific fiber responses and its related consequences to temporal encoding and suprathreshold hearing.

## Conclusions

The current study introduced a refractoriness-masking paradigm to non-invasively isolate AN activity according to the level at which it became effectively recruited using high-fidelity electrophysiological recordings. By utilizing rapid click trains with cumulatively increasing intensities and an inter-click interval of 250 μs, we were able to sequentially mask lower-threshold fibers and isolate AN fiber activity that was newly recruited at higher sound levels. Identifying fiber-specific response patterns helps to bridge the gap between human and animal research in auditory nerve function and may hold potential in developing metrics for peripheral neural degeneration (e.g., cochlear synaptopathy).

## Supporting information

Supplemental Methods and Results

