## Supplemental Methods and Results for "Refractoriness-Based Masking Yields Threshold-Specific Human Auditory Nerve Responses"

#### Behavioral Measures

##### EHF test

The extended high frequency (EHF) test aimed to measure the upper frequency limit of hearing for each participant, following the protocol of Rieke et al. (2017). The target stimulus consisted of a pure tone presented at a fixed intensity of 80 dB SPL. The tone duration was 500 ms, including 20 ms onset and offset ramps to minimize spectral splatter. A continuous background masking noise was presented to the contralateral ear. This noise was generated as a band-limited pink noise (1/*f* power spectrum), spanning frequencies from 4000 Hz to 23999 Hz. The root-mean-square (RMS) amplitude of the masking noise was set 30 dB below that the RMS amplitude of the target tone. Each trial consisted of a 500 ms stimulus presentation followed by a 200 ms inter-stimulus interval.

Each participant completed a total of 6 runs (3 runs per ear). During the test, participants were instructed to press and hold the spacebar when the sound was audible and release it when it was inaudible. The frequency of the target tone was controlled by an adaptive tracking procedure, bounded between 2 kHz and 22.627 kHz. Each run continued until 12 reversals were recorded. The run began at a starting frequency of 4 kHz. The frequency remained constant for the first 4 trials to allow the participant to recognize the stimulus. Initially, the step size was 1/6 of an octave. After the second directional reversal, the step size was reduced to 1/12 of an octave to improve threshold precision.

The frequency threshold for each run was calculated by averaging the frequencies of the last 10 reversals. The frequency threshold for each ear was calculated by averaging the frequency thresholds across the three runs.

##### CRM test

The Coordinate Response Measure (CRM) test followed a protocol previously established by Polonenko and Maddox (2025), based on (Gallun et al., 2013). During the test, participants were required to listen to a target CRM phrase spoken by a male talker with the callsign “Charlie” (e.g., “Ready Charlie, go to blue one now”) embedded in a multi-talker masker. Then, they were asked to report the color (red, white, blue or green) and number (integers from 1 to 8) spoken by the target speaker (such as “blue” and “one” in the above example). Two breaks of a minimum of 15 s were forced during the experiment, and participants were allowed to take other breaks *ad libitum*, including extending the forced breaks.

Before testing, participants first completed a training session in which target phrases were presented at a signal-to-noise ratio (SNR) of 25 dB. After each response, a binomial test was conducted under the null hypothesis that responses were random. Participants were allowed to proceed to the main test session only if the null hypothesis was rejected at an alpha value of 0.01. Depending on performance, the number of trials varied from 5 to 8. If participants failed, the training session was repeated with a 3 dB increase in SNR.

Each trial included a target CRM sentence and a masker sound. The target phrases were drawn from standard CRM sentences (Brungart, 2001). The masker comprised concatenated 10-second segments from 4 distinct male speakers reading audiobooks sourced from the LibriVox public domain collection: *Alice’s Adventures in Wonderland* (Carroll, 2020), *The Adventures of Pinocchio* (Collodi, 2012), *The Time Machine* (Wells, 2011), and *The Wonderful Wizard of Oz* (Baum, 2007). Each masker was truncated to match the duration of the CRM sentence.

Target sentences were presented at 65 dB SPL. The masker level was controlled by an adaptive tracker, with a maximum level of 80 dB SPL. The starting value of the SNR was set at 10 dB. The tracker used a one-up, one-down rule: an initial step size of 5 dB for the first three reversals, followed by a step size of 1 dB for the eight subsequent reversals. Each participant completed three blocks (adaptive tracker). Those with a threshold standard deviations (SDs) ≥4 dB across these blocks were required to complete three additional runs. Only the subsequent valid blocks were used in the analyses.

Thresholds were calculated by averaging the SNRs of correctly answered trials during the last eight reversals in each block. A correct trial was defined as the correct identification of both color and number. Final thresholds were obtained by averaging across the three valid trackers. Given four color choices and eight number options, the chance level for a correct response was 3.1% (1 in 32).

##### AM detection test

The amplitude modulation (AM) detection test was based on the protocol described by Whiteford et al. (2017). During the test, participants listened two sounds—one modulated and one unmodulated—in randomized order. Then, they were asked to identify the AM sound with no time limit. Participants were allowed self-paced breaks after each block.

Before testing, all participants underwent a training session in which the modulation depth was -1 dB. A binomial test was conducted under the null hypothesis that responses were random. Participants were allowed to proceed to the main test session only if the null hypothesis was rejected at the alpha value of 0.01. Depending on performance, the number of trials varied from 5 to 8. If participants failed, the training session was repeated with a 3 dB increase in modulation depth.

Stimuli were generated at a sampling rate of 48,000 Hz. The stimulus length was 2 seconds with 10 ms ramp-up and ramp-down windows. The AM frequency was 5.5 Hz. The AM detection test included four sessions. In two of the sessions, the target stimulus was a 2000 Hz pure tone at 70 or 30 dB SPL for individual sessions. In the other two sessions, the stimuli were modulated click trains. Each click train was a 2-second stimulus consisting of fifty 100 μs click sounds. The sound levels were set at 45 dB or 90 dB peSPL for these individual sessions.

The modulation depth for each session was controlled by an adaptive tracker, ranging from 0 dB (i.e., no modulation) to negative infinity (i.e., fully modulated). The tracker started at –8 dB and followed a one-up, one-down rule with a 2:1 up/down step size ratio. The initial step sizes were 6 dB for the first two reversals, 2 dB for the subsequent two reversals and 1 dB for the final six reversals. Each session included three blocks. Participants with a depth SDs ≥4 dB across their first three blocks completed three additional runs, and only the subsequent valid blocks were used in analyses. The threshold depth of each tracker was calculated by averaging the depth of correct trials obtained during the last six reversals foreach tracker. The final threshold depth was determined by averaging the depth across the three valid trackers.

### Supplementary Results

#### Failure to inhibit fiber-specific responses at the inter-click interval of 1 ms

The *Isolate* response parameters were plotted in **Supplementary Fig. 1B&C.** There was a significant main effect of level on CAP amplitude (*F (4,84) = 5.579, p < .001, η²_p_ = .210*) and latencies (*F (4,84) = 94.156, p < .001, η²_p_ = .818*). We observed a consistent decreasing trend in CAP latency as the sound level increased (*p < .05*), whereas the CAP amplitude reached a plateau as the sound level increased to 90 dB peSPL (*p < .05*). We then tested in the 1 ms ICI sub-session whether preceding lower-level clicks in a click train selectively suppressed lower-threshold AN fibers from responding to subsequent clicks due to the refractoriness, which served as the foundation of this paradigm. The CAP for the *Validate* responses was extracted and compared with those to *Isolate* responses. If lower-threshold fibers were successfully kept in refractory period, the *Validate* responses should be limited regardless of sound levels, compared to that of the *Isolate* responses that should be dominated by fibers that are responsive but not yet saturated. Insufficient CAP waves were observed in the *Validate_45_* and *Validate_60_* conditions, presumably due to low sound levels.

However, we still observed the CAP components in the *Validate_75_* and *Validate_90_* conditions (**Supplementary Fig. 1A)**. As sound levels increased, the CAP and ABR wave V became less synchronized in general. For the CAP amplitude, we observed significant main effect of level (*F (1,13) = 33.495, p < .001, η²_p_ = .347*), and significant main effect of condition (*F (1,13) = 33.379, p < .001, η²_p_ = .102*) but no significant level*condition interaction (*F (1,13) = 4.29, p = .059, η²_p_ = .01*) (**Supplementary Fig. 1D)**. Post-hoc analysis showed that the CAP amplitudes of the *Validate* response were significantly smaller than those of the *Isolate* response, presumably due to refractoriness (*p < .05*). The results were same for the CAP latency (level: *F (1,13) = 23.155, p < .001, η²_p_ = .213;* condition: *F (1,13) = 0.828, p < .001, η²_p_ = .142;* level* condition: *F (1,13) = 1.241, p = .285, η²_p_ = .018*) (**Supplementary Fig. 1E**). While *Validate* responses showed delayed latencies compared to *Isolate* responses at both levels, this was only significant at 90 dB peSPL level (75 dB: *p = .057*; 90 dB: *p < 0.05*). The ABR parameters are shown in **Supplementary Fig. 1F&G** and the results are reported in **Supplementary Table 1**.

The substantial *Validate* responses suggested that the 1ms of inter-click interval might be insufficient to maintain the AN fibers excited by preceding lower-level clicks in a refractory state, leaving them still responsive to subsequent click stimuli. This was also supported by the morphology of the CAP and ABR wave V at high level conditions that showed multiple potential wave components within the window of interest. Therefore, we were unable to disentangle fiber responses to the final click from those contributed by lower-threshold fibers at the 1 ms ICI.

#### No correlation between behavioral measures and neural measures

We conducted a further exploratory analysis to investigate whether any *Isolate* responses could predict any behavioral changes previously proposed to be linked with the neural deficits of higher-threshold fibers in previous human studies, including AM detection thresholds (Çildir et al., 2022; Prendergast et al., 2019), EHF (Motlagh Zadeh et al., 2019) and CRM (Bramhall & McMillan, 2024; Prendergast et al., 2017). The CAP amplitudes and RMS of *Isolate* responses at 75, 90 and 105 dB peSPL in the 250 μs sub-session were correlated with each behavioral measure. However, we did not observe any significant correlation between any neural and behavioral measures at any level (**Supplementary Fig. 2**).

Since the amplitude of CAP can be influenced by non-sensory factors, such as the placement of the TM electrode or individual differences of eardrums, we decided to minimize these potential effects on CAP amplitudes by calculating the amplitude ratio of two consecutive *Isolate* conditions. If isolated responses represent fibers with different response thresholds, the ratio of isolated responses might serve as an indicator of AN fiber recruitment. Yet there was no significant correlation between any amplitude ratio and any behavioral measures (**Supplementary** **Fig. 3**).  Nevertheless, it should be noted that the participants recruited in this study were young listeners with normal hearing thresholds and no noise exposure history was collected. Future studies should recruit participants who struggle with speech perception in noise to verify the sensitivity of the neural measures in predicting perceptual deficits.

Supplementary **Table 1.** ANOVA of ABR parameters in the 1 ms experimental sub-session.

| **Parameters** | **Source** | ***df*** | ***F*** | ***p-value*** | ***η²_p_*** |
| --- | --- | --- | --- | --- | --- |
| Amplitude | Level | 1 | *0.218* | *.657* | *.011* |
|  | Condition | 1 | *0.883* | *.384* | *.015* |
|  | Level* Condition | 1 | 11.984 | .013* | .209 |
|  | Residual | 6 |  |  |  |
| Latency | Level | 1 | *0.543* | *.489* | .014 |
|  | Condition | 1 | *8.391* | *.027** | *.136* |
|  | Level* Condition | 1 | 0.528 | .495 | .025 |
|  | Residual | 6 |  |  |  |

*Note: *p < .05*

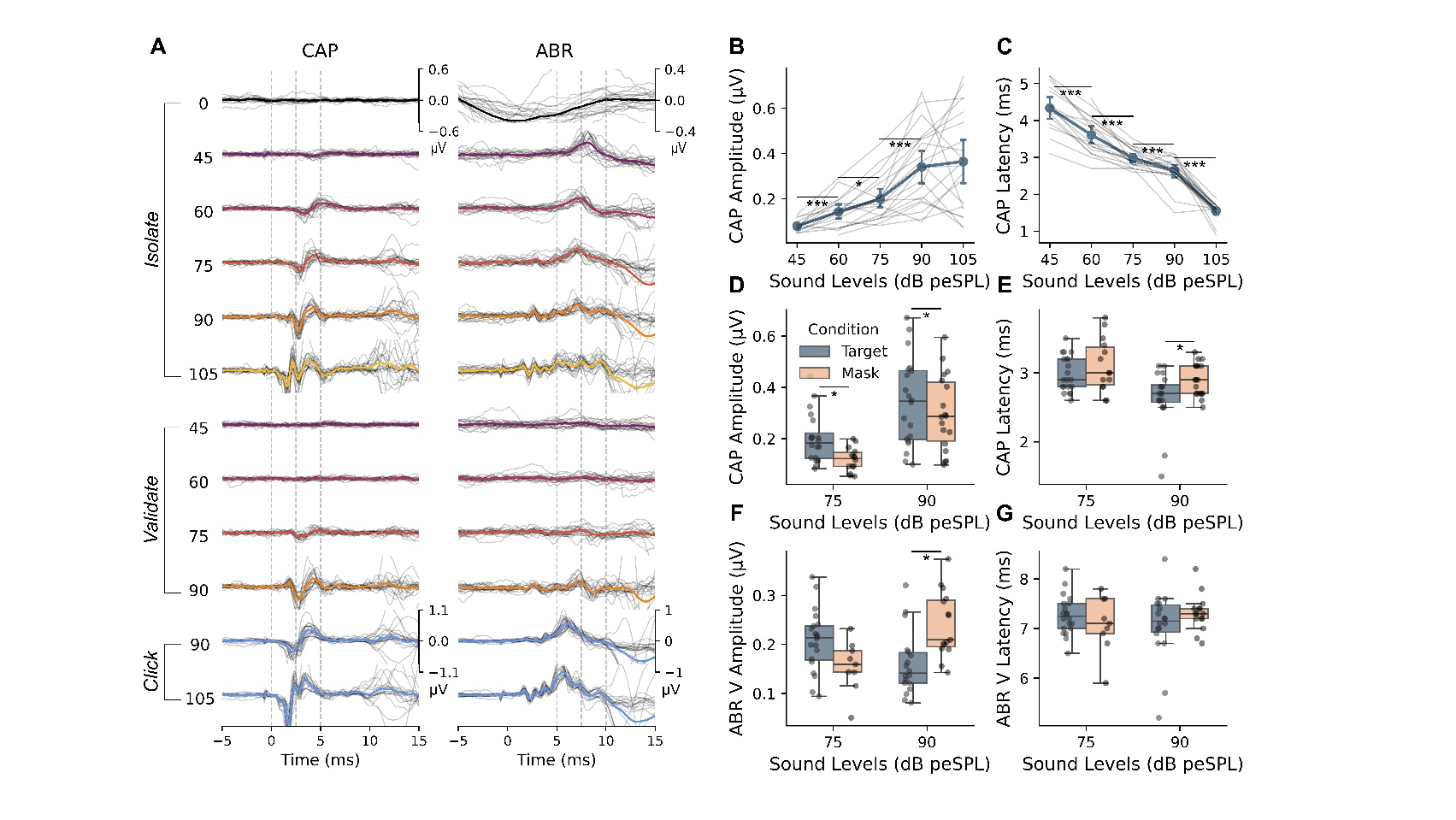

**Supplementary Fig. 1** Grand average CAP and ABR waveforms in the 1 ms experimental sub-session. (A) Grand average CAP and ABR waveforms showed that substantial CAP components remain in the *Validate_75_* and *Validate_90_* conditions, indicating a failure to fully inhibit fiber-specific responses with a 1 ms inter-click interval. (B-C) *Isolate* CAP amplitudes and latencies across increasing sound levels, showing a consistent decreasing trend in latency and an amplitude plateau starting at 90 dB peSPL. (D-E) Comparisons of CAP amplitudes and latencies between *Validate* and *Isolate* conditions at 75 and 90 dB peSPL, suggesting that *Validate* responses have significantly smaller amplitudes and delayed latencies compared to *Isolate* responses. (F-G) Corresponding ABR wave V amplitudes and latencies for the *Isolate* and *Validate* conditions.

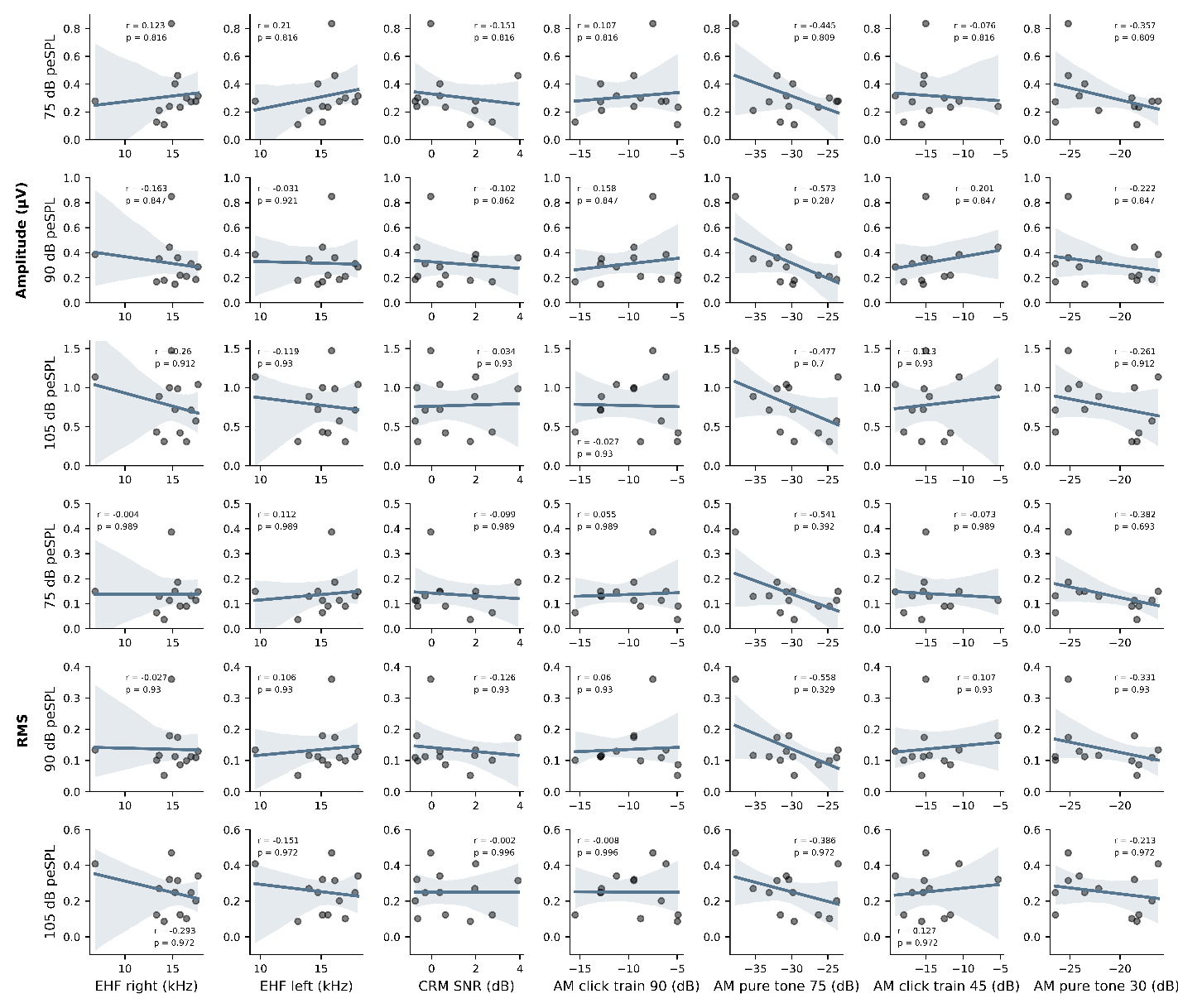

**Supplementary Fig. 2** Correlation the behavioral and *Isolate* response parameters in the 250 μs experimental sub-session. Scatter plots illustrating the relationship between two neural response parameters (CAP amplitudes and RMS values of *Isolate* responses at 75, 90, and 105 dB peSPL) and the behavioral assessments. The behavioral tasks evaluated include the Extended High Frequency (EHF) test, the Coordinate Response Measure (CRM) test for speech-in-noise perception, and the Amplitude Modulation (AM) detection test. No significant correlations were observed between any neural parameters and behavioral measures at any sound level.

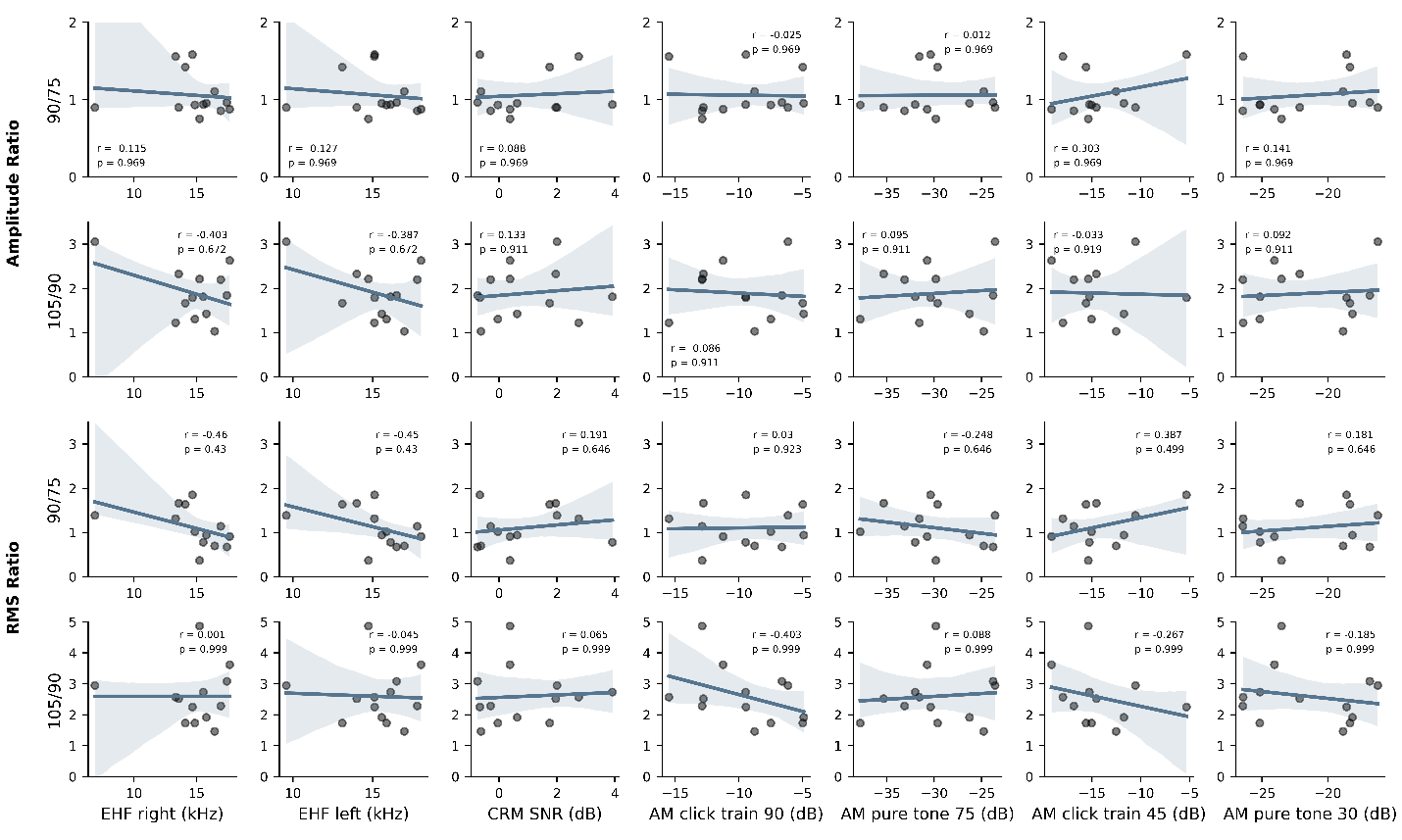

**Supplementary** **Fig. 3** Correlation the behavioral and neural response ratio in the 250 μs experimental sub-session. Scatter plots showed no significant relationship between all behavioral measures (EHF, CRM, and AM detection thresholds) and the CAP amplitude and RMS ratios of two consecutive *Isolate* conditions. The ratios were calculated to normalize the data and minimize the influence of non-sensory factors, such as the physical placement of the tympanic membrane electrode.
